# Comparison of Fixed Single Cell RNA-seq Methods to Enable Transcriptome Profiling of Neutrophils in Clinical Samples

**DOI:** 10.1101/2024.08.14.607767

**Authors:** Klas Hatje, Kim Schneider, Sabrina Danilin, Fabian Koechl, Nicolas Giroud, Laurent Juglair, Daniel Marbach, Philip Knuckles, Tobias Bergauer, Matteo Metruccio, Alba Garrido, Jitao David Zhang, Marc Sultan, Emma Bell

**Affiliations:** Roche Pharma Research and Early Development, Pharmaceutical Sciences, Roche Innovation Center Basel, F. Hoffmann-La Roche Ltd, Grenzacherstrasse 124, 4070 Basel, Switzerland

**Keywords:** Neutrophils, single cell RNA-seq, transcriptome, biomarker

## Abstract

Monitoring neutrophil gene expression is a powerful tool for understanding disease mechanisms, developing new diagnostics, therapies and optimizing clinical trials. Neutrophils are sensitive to the processing, storage and transportation steps that are involved in clinical sample analysis. This study is the first to evaluate the capabilities of technologies from 10X Genomics, PARSE Biosciences, and HIVE (Honeycomb Biotechnologies) to generate high-quality RNA data from human blood-derived neutrophils. Our comparative analysis shows that all methods produced high quality data, importantly capturing the transcriptomes of neutrophils. 10X FLEX cell populations in particular showed a close concordance with the flow cytometry data. Here, we establish a reliable single-cell RNA sequencing workflow for neutrophils in clinical trials: we offer guidelines on sample collection to preserve RNA quality and demonstrate how each method performs in capturing sensitive cell populations in clinical practice.

## Introduction

Neutrophils are innate immune effector cells that comprise approximately 60% of leukocytes in circulation. They mediate the body’s first response to invading microorganisms through degranulation, phagocytosis and the production of Neutrophil Extracellular Traps (NETs)(Papayannopoulos, 2018). Neutrophil dysregulation, particularly NET formation, is strongly implicated in human diseases ranging from sepsis, autoimmunity to cancer metastasis and inflammatory diseases (Papayannopoulos, 2018). In the clinic, neutrophils and neutrophil expression signatures are increasingly being used as biomarkers. Notably, the neutrophil-to-lymphocytes ratio, when combined with tumor mutation burden, is being used to forecast the effectiveness of immune checkpoint inhibitors in cancer treatment (Salcher et al., 2022; Valero et al., 2021). Additionally, biomarkers derived from neutrophils are being investigated for their potential to predict major adverse cardiac events (Yiu et al., 2023).

Single cell sequencing has helped to improve our understanding of the different transcriptional states of neutrophils, and suggested a future role for the neutrophil gene expression signatures as clinical biomarkers. Four distinct and stable transcriptomic states observed during the maturation and activation of neutrophils have been described: Nh0, Nh1, Nh2 and Nh3 (Wigerblad et al., 2022) suggesting that a deeper understanding of these transcriptomes could provide disease biomarkers. Expanding on this (Montaldo et al., 2022) have described the transcriptome of neutrophils in a steady state and upon stress using both bulk RNA-seq approach and scRNA-seq on live cells using 10X 3 prime library methods that were modified to capture neutrophils. Here they describe how the different activation status of neutrophils are predictive biomarkers for organ transplant success (Montaldo et al., 2022). A recent study has highlighted how there is a high level of transcriptional heterogeneity in neutrophils isolated from different cancer types. Originally, high levels of invading neutrophils were thought to be a poor prognostic indicator. However, more recent findings suggest that neutrophils with an antigen presenting transcriptional program are associated with a positive outcome in most cancers (Wu et al., 2024b). Understanding of neutrophil biology and phenotypes will help develop biomarkers for identifying patients that may experience cytokine release syndrome in response to T-cell engaging therapies, for example T-cell bispecifics (Leclercq et al., 2022). Taken together, these papers suggest a future requirement for profiling neutrophils in clinical samples.

Neutrophils contain lower RNA levels than other cell types in the blood (Wigerblad et al., 2022). Classical methods using gel emulsion beads (e.g. 10X) have proved challenging to capture neutrophils and granulocytes (Salcher et al., 2022). Indeed, without modification, 10X 3 prime transcriptomic methods are unable to capture the transcriptomic profiles of neutrophils. However, several groups have demonstrated that generating neutrophil scRNA-seq data is technically feasible, even if there is a high percentage of loss. (Wigerblad et al., 2022) detailed a method where addition of an RNAse inhibitor and modifications to the bioinformatic pipeline was sufficient to capture the transcriptome of neutrophils. As part of earlier work looking at neutrophils in whole blood, we showed that the microwell based scRNA-seq BD Rhapsody effectively captured the transcriptome from neutrophils. The percentage of neutrophils retrieved from samples was comparable to results from flow cytometry using CD16, CD11b and CD62L as markers (Leclercq et al., 2022). A direct comparison between the BD Rhapsody and 10X 3 prime suggested that RNA capture is significantly more sensitive in the microwell based method, leading to more sensitive detection of cells with a low RNA content (Salcher et al., 2024).

Single cell RNA-seq is a powerful tool in drug discovery. However, its potential for use with clinical samples is limited by the requirement to use fresh cells. For PBMCs, protocols have been developed whereby cell separations can be performed at the clinical sites and then cells cryopreserved and banked for later analysis at a central testing facility (van der Wijst et al., 2018). However, for more sensitive cell types such as neutrophils, this is not possible, as a high proportion of these cells die and the remaining cells are morphologically and functionally altered in the freeze-thaw process (Braudeau et al., 2021; de Ruiter et al., 2018; Verschoor et al., 2018). For these cell types, the single cell analysis must be performed at the clinical site, which reduces the number of clinical sites that are able to collect samples for scRNA-seq analysis. Taken together the biological importance, sensitive nature of neutrophils in combination with the complexity of global clinical trial settings call for an easy-to-use stabilization protocol for subsequent single cell RNAseq.

We selected three new technologies to compare: Evercode from PARSE technologies, 10X Genomics FLEX solution, and the Honeycomb Technologies HIVE device. The selection criteria was based on the ability to stabilize cells rapidly prior to library prep, the requirement to process large number of cells and a commercially available product that can be distributed easily to clinical sites. PARSE technologies scRNA-seq works on a principle of combinatorial barcoding, where fixed cells are given a sample barcode with the reverse transcription step, samples are then pooled and split before a further three successive barcoding steps, including the addition of a unique molecular barcode (Rosenberg et al., 2018). This approach allows for up to 96 multiplexed samples, and has been reported to detect more genes expressed at low levels than the 10X 3 prime library prep (Xie et al., 2020). The HIVE device works on the principle that cells are distributed into nano-wells and stabilized. The samples can be stored at -80°C prior to the library preparation steps. The HIVE device has successfully been used to isolate neutrophils from RBC-depleted donor samples (Sheerin et al., 2023). In the 10X RNA Flex fixed and permeabilized cells are incubated with a set of 18,532 probes covering the entire transcriptome prior to library preparation steps. The use of probe hybridization allows for the capture of smaller fragments of RNA which are found in formalin fixed, paraffin embedded tissue. This method has been successfully used on FFPE tissues and xenograft models (Llora-Batlle et al., 2024; Wang et al., 2023).

Neutrophils are reported to have a short half-life ex vivo and the methods of isolation can lead to activation or apoptosis. Therefore, we used the findings of previous studies on neutrophil isolation to define the conditions for this study. Previous reports have demonstrated that neutrophils suitable for functional characterization can be isolated from blood stored at room temperature or at 4°C for 24 hours, or up to 72 hours when stored at 37°C (Bonilla et al., 2020; Li et al., 2024; Wood et al., 1999). Incubators for sample storage are not always be available at clinical sites, therefore we opted to look at the impact of storage at 4°C for 24 hours. Currently, there is little information exploring the effect of time from blood draw to analysis or fixation on neutrophil transcriptome stability. This work aims to evaluate the new generation of fixed single cell technologies to determine their suitability for 1) measuring the neutrophil transcriptome and 2) their potential for implementation in clinical trials which require minimal sample processing and sample stabilization.

## Results

### Study design

To compare the different technologies, blood was drawn from healthy donors and then divided into different aliquots which were tested using 10X Flex, PARSE, and 10X 3 prime chemistries (Figure 1A). An aliquot for each donor was run on the flow cytometer to characterize cells into the major cell types to compare with the results from the scRNA-seq clustering. We evaluated the HIVE devices in a separate experiment using the same format: The blood samples were profiled using HIVE, 10X 3 prime and flow cytometry in parallel (Figure 1B). In order to compare directly across the technologies we limited our analysis to the 18,532 genes captured in the 10X RNA FLEX probe set. We used our established BESCA pipeline (Madler et al., 2021). The knee plots (Supplementary Figure 1) reveal a clear separation between cells and empty droplets for PBMC isolation, aiding in cutoff determination. However, RBC-depleted samples lack this distinct separation due to low gene expression in granulocytes. To ensure inclusion of neutrophils, we applied a minimum threshold of 200 genes and 200 UMIs across all samples.

**Figure 1:**
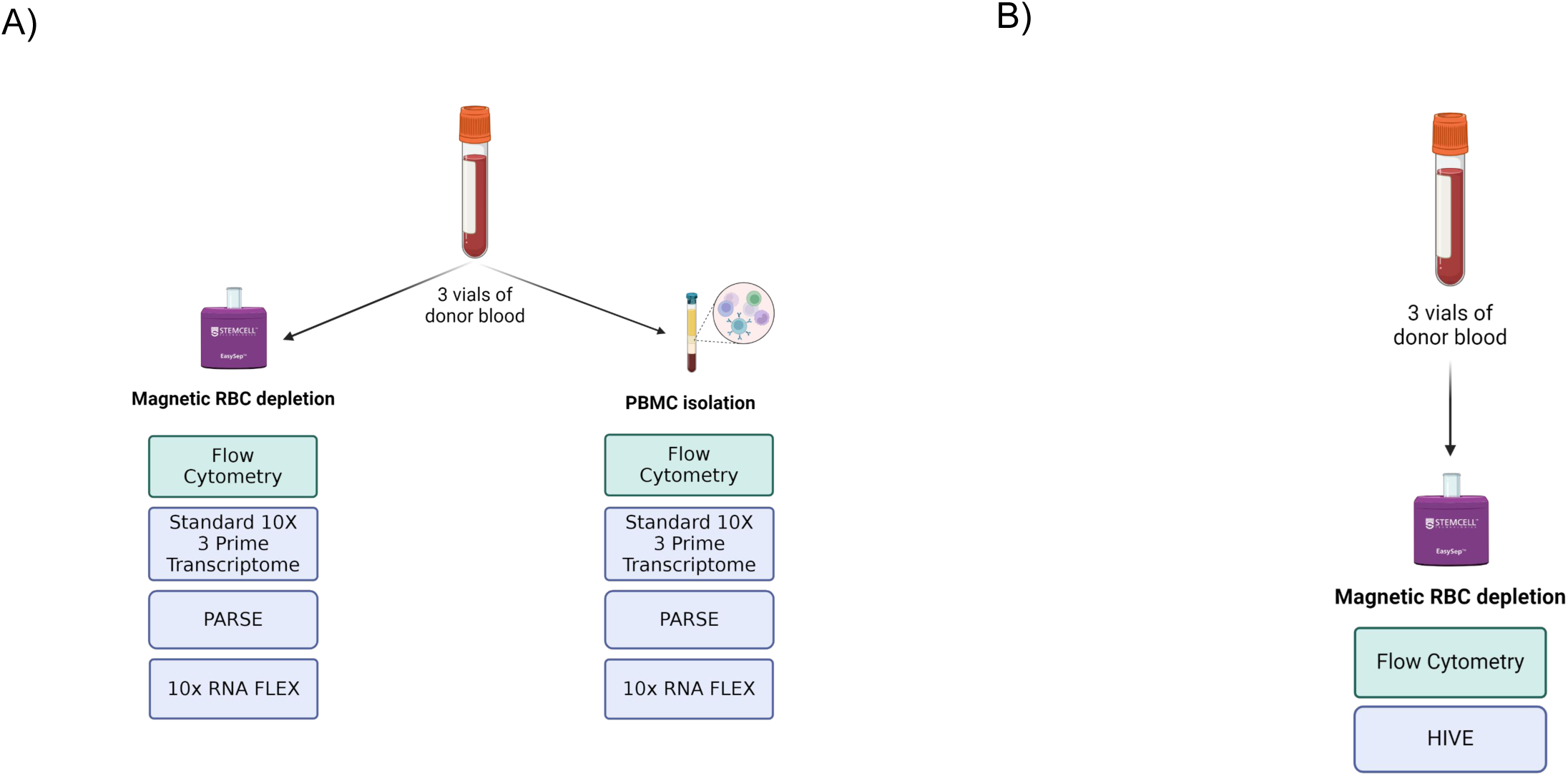
Diagram showing the experimental design to test the four different technologies. (A) PARSE, 10x FLEX, 10x 3 prime and flow cytometry were tested on the same three blood samples, and (B) HIVE was tested on a different set of blood samples from three different donors at a different date. The samples were also profiled using flow cytometry.

### Comparing the quality of scRNA-seq from the different methods

We compared the quality of the scRNA-seq data using the following parameters: UMI counts, the number of genes detected and the percentage of mitochondrial genes (Figure 2A). These parameters are used to discriminate low quality cells where the cells are stressed, or cell leakage occurring during processing (Ilicic et al., 2016). Across all of the scRNA-seq technologies the mitochondrial gene expression levels were low, between 0-8%, with PARSE showing the lowest levels of mitochondrial gene expression, followed by 10X RNA FLEX. 10X 3 prime samples and HIVE which used non-fixed cells as input both had higher levels of mitochondrial genes detected. For all the novel methodologies, the number of genes detected and the number of UMIs were lower in the RBC depleted samples compared to the PBMC samples (Figure 2B & C, Supplementary Figure 2). For the RBC depleted samples, we observed a bimodal distribution in the violin plots. This was due to two populations of cells with different overall gene expression levels: the PBMC population with high gene expression and the granulocytes with low levels of genes expressed per cell and lower mRNA levels in general (Wigerblad et al., 2022).

**Figure 2:**
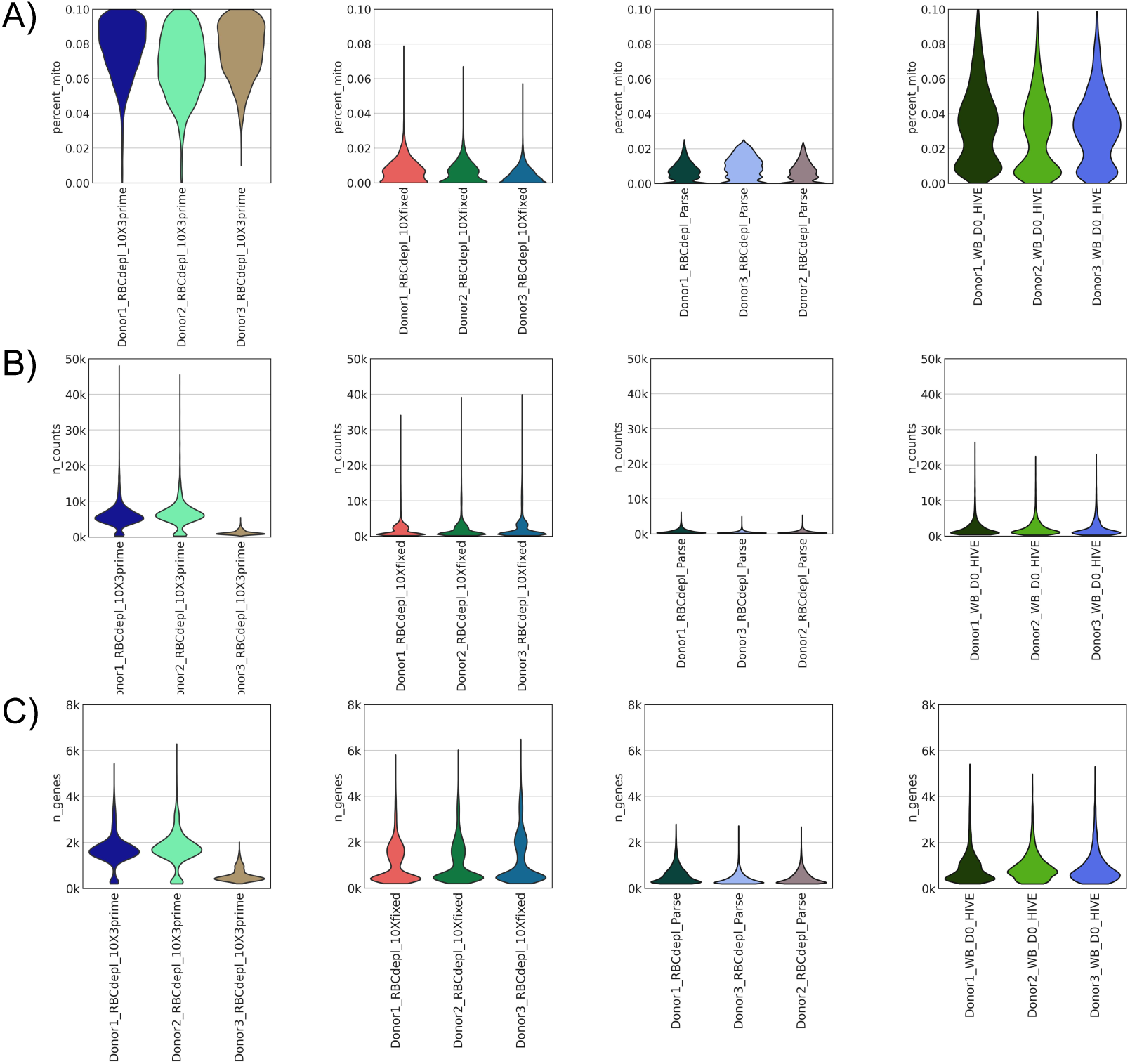
Violin plots showing the (A) mitochondrial gene expression levels (B) n_counts and (c) UMI counts for 10x 3 prime, 10X FLEX, PARSE and HIVE for RBC depleted samples.

Next, we examined the dynamic range of 10X Flex, Parse, and HIVE alongside the 10X 3 prime technology. To do this we examined the expression of genes that have been classified with high (B2M), high-medium (ACTB), low-medium (CTCF) and low expressed (PGK1). In 10X Flex, HIVE and 10X 3 prime we observed that the majority of the cells were expressing high levels of B2M and ACTB, with the number of cells expressing PGK1 and CTCF reducing and the magnitude of expression also decreasing (Supplementary Figures 3 &4). For the PARSE PBMC samples, the number of counts per cell were comparable with the 10X 3 prime and 10X Flex samples. However, the PARSE samples still showed lower expression of B2M and ACTB, and higher levels of the lower expressed genes PGK1 and CTCF. From this data, we conclude that the PARSE samples show a different dynamic range to the 10X data and HIVE data with a greater representation of genes with lower levels of expression.

### scRNA-seq Clustering

We combined the data from the different technologies for clustering purposes and observed that the cells clustered based on the technology used (Figure 3A). Within the technology-specific clusters we observed that the cells separated into clusters based on the cell separation method used (PBMC versus RBC depletion) (Figure 3A), and finally the cells clustered into different cell types (Figure 3A). We observed that the neutrophils clusters in all technologies were associated with lower n_counts and UMI counts which is in line with the low levels of RNA and gene expression in this cell population (Figure 3B). The 10X 3 prime and HIVE clusters showed higher percentage mitochondrial gene expression (Figure 1A and 3B). Looking at the cell type clustering for each individual technology (Figure 3A & Supplementary Figure 5), we see that the major cell types can be identified clearly in the four different technologies, however neutrophil clusters were absent in the 10x 3 prime. The cell types separated into more defined clusters for both the 10X technologies. In all technologies, we also observed artifact clusters, which are composed of empty droplets due to the lower cutoffs used, doublets or cell types that cannot be assigned to any group. The throughput of the PARSE and 10X FLEX technologies was much higher than the 10X 3 prime and HIVE technologies. We did observe a high level of doublets in the 10X Flex RBC-depleted samples (∼20%) compared to the other technologies. However, we could easily identify the doublets and could remove them from the analysis (Figure 3A & Supplementary Figure 3C).

**Figure 3:**
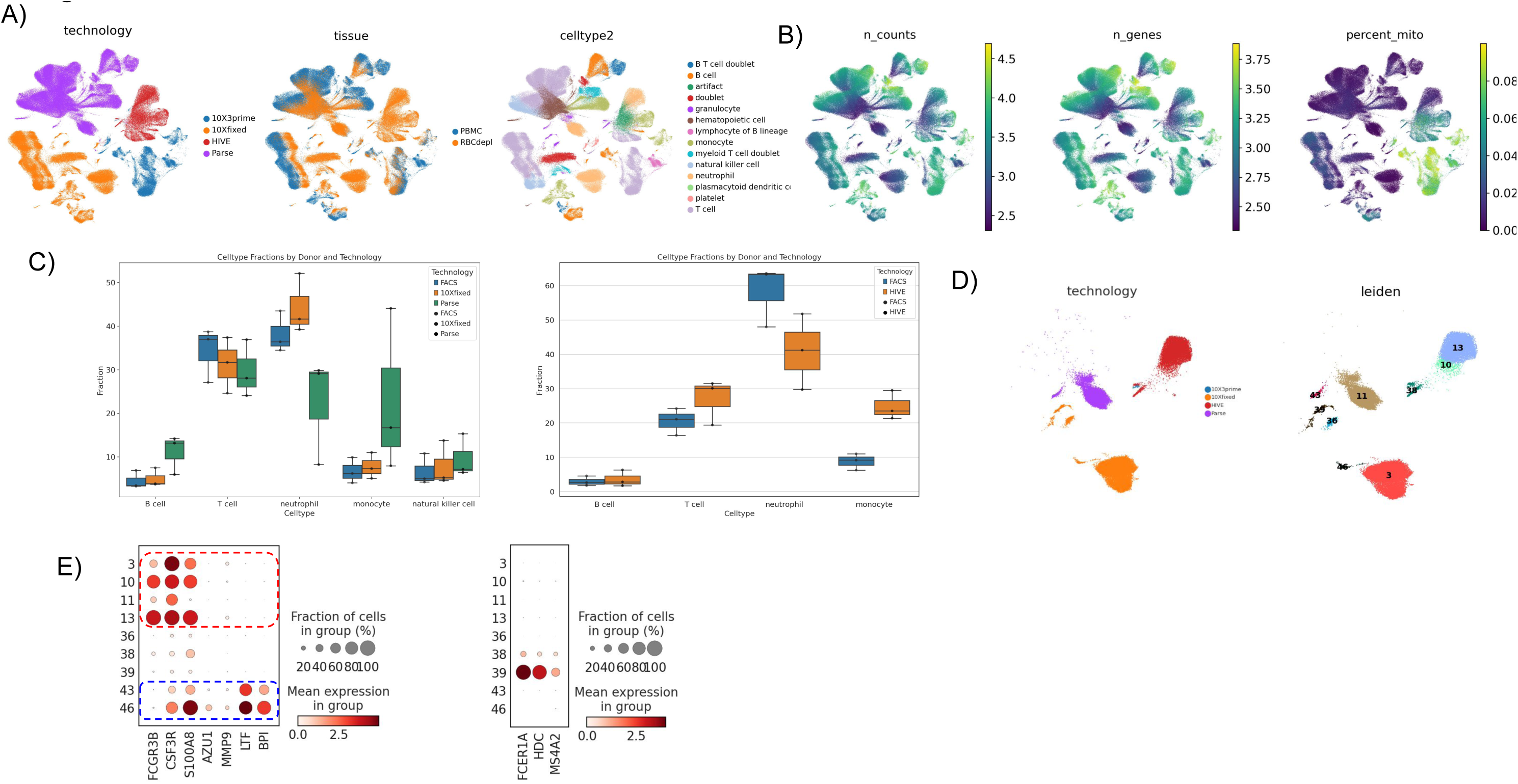
UMAPs cell separations with color coding denoting (A) technology, cell separation used and % different cell types. (B) The quality control parameters are shown on UMAPs color coded by UMI counts, n_counts and mitochondrial gene expression. (C) Box plots showing the % of B cells, T cells, neutrophils, monocytes and NK cells determined by flow cytometry, FLEX and PARSE or flow cytometry and HIVE. (D) UMAP of the granulocyte clusters coloured by technology, unsupervised clustering of granulocytes. (E) Dot plot showing the expression of the top 5 marker genes per cluster, for those clusters with >100 cells.

### Percentage cell populations determined by scRNA-seq compared to flow cytometry

In order to determine how well the fixed cell scRNA-seq technologies captured neutrophils we compared the percentage neutrophils from each technology with the percentage neutrophils determined by flow cytometry on the same sample. Please note that for the 10X RNA FLEX, 10X 3 prime and PARSE blood for the same 3 donors was tested. For the HIVE evaluation, blood from a different three was tested. We profiled an aliquot from each blood sample by flow cytometry to identify different cell types. 10X FLEX, PARSE and HIVE all successfully isolated neutrophils from the red blood cell depleted samples. The percentage of neutrophil populations using FLEX were the closest to those determined by flow cytometry (Table 2, Figure 3C, Supplementary Tables 1, 2, 3, 4, 5 & 6; Supplementary Figure 6). FLEX and PARSE also compared favorably with flow cytometry results for the isolation of T-cells, B-cells, Monocytes, Natural Killer cells. Indeed, the performance was comparable on the PBMC isolations with the 10x 3 prime methods and flow cytometry (Supplementary Tables 1, 2, 3, 4, 5 & 6).

### Identification of neutrophil populations

We looked at the clustering within the granulocyte clusters for the different technologies to determine if different populations of neutrophils could be identified. Unsupervised clustering demonstrated that two different populations of neutrophils can be defined in the FLEX (Cluster 46) and PARSE (Cluster 43) data (Figure 3D), these clusters have elevated expression of LTF and BPI compared to the other granulocyte clusters. A second group consisting of clusters 3 (10X Flex), 10 and 13 (HIVE) is characterized by expression of only FCGR3B, CSF3R and S100A8, the canonical markers for neutrophils found in the blood (Figure 3D). A population expressing Basophil markers (FCER1A, HDC and MS4A2) was defined only in the FLEX data set (Cluster 39) (Figure 3D&E). We were unable to identify Eosinophils in any of the data sets.

### Time course for optimum sampling of neutrophils

Neutrophils are a particularly sensitive cell type with a reported short half-life In Vivo and In Vitro (Lahoz-Beneytez et al., 2016; Scheel-Toellner et al., 2004). In order to determine the maximum time that samples could be stored prior to processing, we tested cells at different time points after blood draw (immediate processing, 2, 4, 6, 8 and 24 hours after blood draw), prior to fixing and measuring transcriptome by 10X Flex. After the cell isolation steps we performed a cell count prior to stabilization (data not shown). We observed little overall cell death or decrease in cell count over the 24 hours after the blood draw. In concordance with this, the general quality of the scRNA-seq data was unchanged across the time course. There was little increase in expression of mitochondrial genes, with all samples having expression levels of <1% for mitochondrial genes, number of counts or UMI counts across the time course (Supplementary Figure 7A, B &C). There were also no differences in the % cell types over time since blood draw as determined the scRNA-seq (Supplementary Figure 7D, E & F), indicating that there is no apoptosis of specific cell types taking place over the 24 hours.

We compared the transcriptional profile of neutrophils at different time intervals after the blood draw and we did observe that the number of genes differentially regulated compared to the 0h time point started to significantly increase 4 hours post blood draw, with the number of genes up and down regulated increasing at each time point (Figure 4A & B). The most significantly changed pathway was associated with cell stress defined as Stress MP5 by (Gavish et al., 2023). In our results, we observed that stress signatures were upregulated in all time points 2 hours after blood draw, showing the importance of prompt sample processing (Figure 4C). This data is concordance with the previous report by (Connelly et al., 2022) which found that markers of neutrophil activation, apoptosis and degranulation 4 hours post blood draw. Therefore, despite live, functionally active neutrophils being present in blood samples 24 hours post blood draw, the transcriptome of neutrophils is significantly changed after 4 hours post blood draw. Our results indicate that immediate fixation of neutrophils is required if the transcriptome is being analyzed.

**Figure 4:**
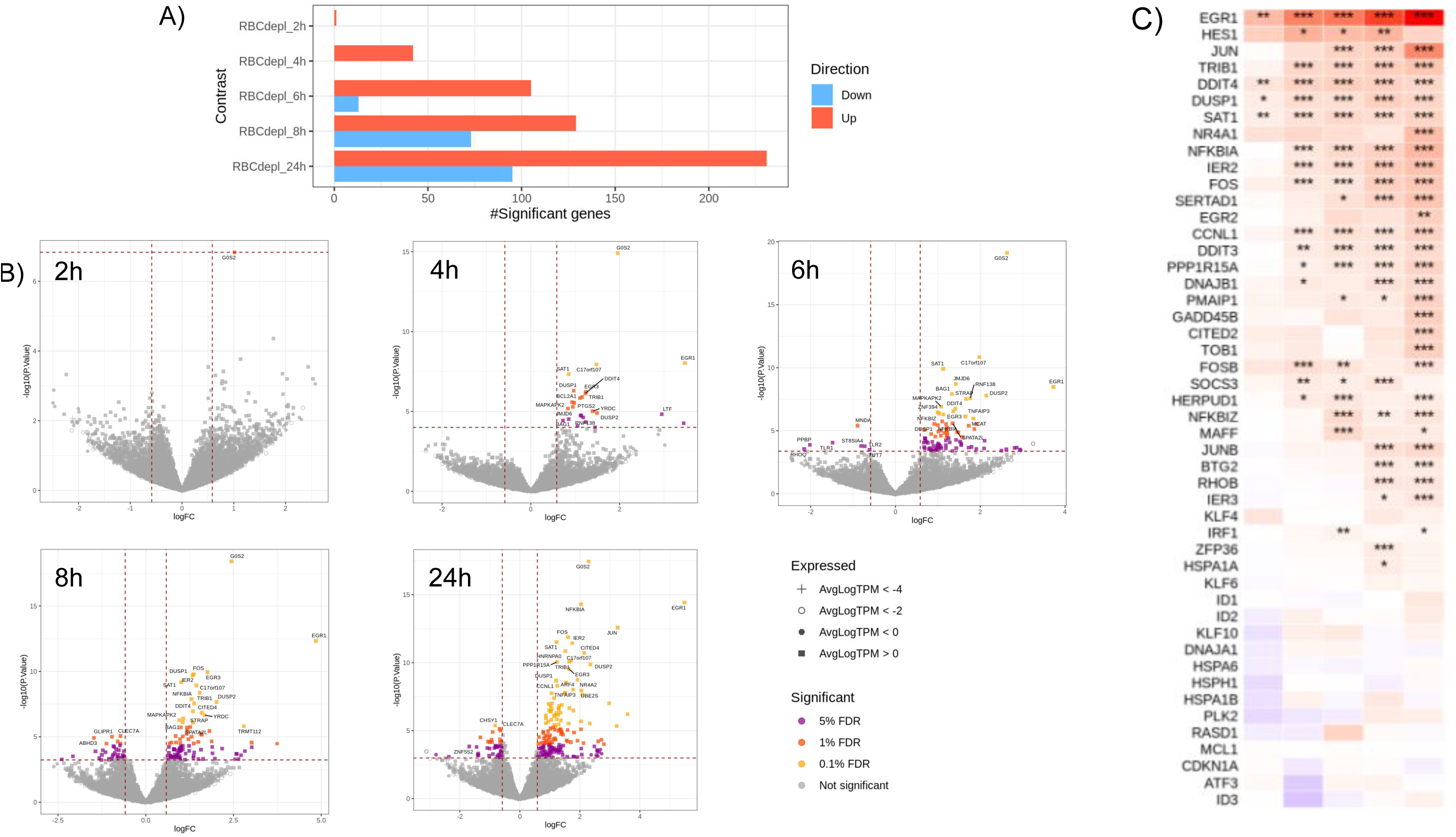
Transcriptional profile of neutrophils after blood draw (A) Graph showing the number of genes differentially regulated (up or down) in neutrophil pseudobulks at 2h, 4h, 6h, 8h or 24h after blood draw compared to the sample processed immediately after blood draw (0h). The data shows results for the 3 donors (n=3) for each time point. (B) Volcano plots for the neutrophil pseudobulk showing magnitude of the genes up or down regulated compared to 0h for each time point tested. (C) Pathway enrichment for cell stress for neutrophil pseudobulks over the time course.

## Conclusions

The recent advances in fixed cell scRNA-seq allowing the rapid stabilization and storage of cells prior to library prep will enable the wider implementation of scRNA seq analysis in clinical trials. 10X FLEX, PARSE and HIVE protocols would support a model where cells are stabilized at the clinical site allowing storage and transport to the analytical labs where library prep and sequencing can take place. Practically, the HIVE devices presents a straightforward protocol for use at a clinical site, with the cells simply pipetted into the device after the cell separation step. We also found that cells stored in HIVE devices at -80C for up to 3 weeks showed good quality data comparable to cells processed immediately (data not shown). For Flex, the samples need to be centrifuged after the cell separation and then resuspended in paraformaldehyde. Since these experiments were completed, 10X have modified their protocols allowing whole blood to be stabilized with paraformaldehyde, then stored and transported at -80°C. This would allow the cell separation and analysis to be performed at the analytical site, presenting a simple procedure for cell stabilization at the clinical site. The PARSE Technologies protocol for fixation of cells requires several consecutive centrifugation steps, making it the protocol that requires the longest hands on time for the fixation steps and practically the hardest to perform at a clinical site.

Our experiments indicate that all three of the technologies produce high scRNA-seq quality data, with low levels of mitochondrial gene expression. FLEX and PARSE, which use fixed cells, have lower levels of mitochondrial gene expression than 10X 3 prime and HIVE that use live or frozen cells as input. This may be due to the release of cytoplasmic RNA on fixation or permeabilization of the cell (De Simone et al., 2024). Interestingly, for PARSE data we observed a different dynamic range than observed for the other technologies. Here we observed that the fully combinatorial barcoding approach (Rosenberg et al., 2018) led to greater sensitivity of detection of genes with lower expression levels. Interestingly, for approaches where combinatorial barcoding techniques were combined with droplet based fluidic systems for scRNA-seq have been reported to have lower sensitivity of detection for genes that have a low level of gene expression in comparison to 10X 3 prime, this maybe due to the combinatorial barcoding being performed inside the droplet, as opposed to within the fixed cell (PARSE) (Datlinger et al., 2021; Wu et al., 2024a). All three fixed scRNA-seq methods successfully captured neutrophils from RBC depleted samples. Our analysis suggested that 10x FLEX captured the different white blood cell components in the red blood cell depleted samples to a similar percentage as flow cytometry. HIVE and PARSE, while capturing neutrophil profiles did not compare as favorably with the flow cytometry results. For cell type assignment in general, we observed the closest alignment with flow cytometry using FLEX, which performed well across all cell types. PARSE and HIVE both had greater deviation from the cell type proportions estimated by flow cytometry. Although technical optimization of the methods and bioinformatic pipeline may improve the cell assignment, these results are in alignment with a previously published study on PBMC which showed that cell type calling for FLEX was closely aligned with CyTOF for the same sample, whereas greater differences were observed with PARSE and HIVE (De Simone et al., 2024).

Unsupervised clustering of the granulocyte cells across all the methods tested defined two distinct groups of neutrophils. The first subtype, expressing BPI and LTF was detected in the FLEX and PARSE data aligns with immature neutrophils: LTF in particular has been shown to be highly expressed at earlier time points in neutrophil differentiation (Grieshaber-Bouyer et al., 2021). A neutrophil type was observed in all three technologies and is characterized by expression of FCGR3B, CSF3R and S100A8 markers of mature neutrophils (Grieshaber-Bouyer et al., 2021). In the FLEX data only, we were able to identify a population of basophils, a rare and sensitive cell type that is <1% of cells in peripheral blood (Min et al., 2012). The cell type was not detected by the other technologies. Taken together, we could only differentiate all three different cell types detected in the 10X FLEX data. FLEX has previously been shown to have more stable gene expression and improved variance compared to HIVE and PARSE (De Simone et al., 2024), we propose that this may play a role in detecting these rare cell types.

This work compared three novel methods to determine if scRNA-seq could be implemented at clinical sites for profiling neutrophils. All three methods produced high quality data and were able to capture neutrophils from peripheral blood samples. However, for clinical samples we determined that FLEX had the best performance, with the proportions of neutrophils captured in blood samples comparable to those observed by flow cytometry and the workflow being the most amenable to sample collection at the clinical site. Additionally using FLEX, we were able to define two distinct populations of neutrophils: immature neutrophils and those expressing canonical neutrophil markers. We recommend that the time between blood draw and fixation is limited to 2 hours, as after this time we observe an increase in differential gene expression regulation associated with stress. To our knowledge, this is the first study to present a route to scRNA-seq implementation in clinical trials and a powerful tool for biomarker development and understanding of neutrophil biology.

## Acknowledgements

We would like to acknowledge the Flow 360 Labs team at Roche for their support in Flow cytometry data acquisition.

## Author Contributions

**KH:** Experimental design, Data Analysis and Interpretation, Manuscript Preparation. **KS:** Experimental design, Data Analysis and Interpretation, Manuscript Preparation **SD:** Experimental Design, Experimental Execution, Manuscript Preparation **FK:** Experimental Design, Experimental Execution, Manuscript Preparation **NG:** Experimental Design, Experimental Execution **LJ:** Experimental Execution, Data Analysis and Interpretation, Manuscript Preparation **DM:** Data Analysis and Interpretation, Manuscript Preparation **PK:** Data Analysis and Interpretation, Manuscript Preparation. **MM:** Experimental design, Manuscript Preparation **TB:** Experimental Design, Manuscript Preparation **AG:** Manuscript Preparation **JDZ:** Data Analysis and Interpretation **MS:** Manuscript preparation **EB:** Experimental Design, Experimental Execution, Data Analysis and Interpretation, Manuscript Preparation.

## Declaration of AI and AI Assisted Technology

During the preparation of this work the authors did not use any AI or AI assisted technology.

## Supplemental Information titles and legends

**Supplementary Table 1:** Comparison of the technologies used in this study

**Supplementary Table 2:** Cell population comparison. Table shows the mean % neutrophils of the three donors tested determined by cell type analysis. For each technology there is a Mean absolute error (MAE) and Root mean squared error (RMSE) which shows the difference between the flow cytometry values and the scRNA-seq derived results.

**Supplementary Table 3:** Donor 1 cell population comparison for Flow cytometry, 10X 3 prime transcriptome, 10X FLEX, PARSE for PBMC isolation and RBC depletion

**Supplementary Table 4:** Donor 2 cell population comparison for Flow cytometry, 10X 3 prime transcriptome, 10X FLEX, PARSE for PBMC isolation and RBC depletion

**Supplementary Table 5:** Donor 3 cell population comparison for Flow cytometry, 10X 3 prime transcriptome, 10X FLEX, PARSE for PBMC isolation and RBC depletion

**Supplementary Table 6:** Donor 4 cell population comparison for Flow cytometry,10X 3 prime transcriptome and HIVE for PBMC isolation and RBC depletion

**Supplementary Table 7:** Donor 5 cell population comparison for Flow cytometry,10X 3 prime transcriptome and HIVE for PBMC isolation and RBC depletion

**Supplementary Table 8:** Donor 6 cell population comparison for Flow cytometry,10X 3 prime transcriptome and HIVE for PBMC isolation and RBC depletion

**Supplementary Table 9:** Flow cytometry antibodies

**Supplementary Figure 1:** Knee plots for PARSE, FLEX and HIVE data for number of genes expressed per cell and UMI counts per cell for (A) PBMC and (B) RBC depleted samples.

**Supplementary Figure 2:** Violin plots showing the (A) mitochondrial gene expression levels (B) n_counts and (C) UMI counts for 10x 3 prime, 10X FLEX and PARSE for PBMC samples.

**Supplementary Figure 3:** A comparison of the dynamic range for the 10X 3prime, 10X FLEX and PARSE for 4 different genes that represent highly expressed (B2M), medium high expression (ACTB), medium low (PGK1) and low (CTCF) expression. These graphs show the expression in the RBC depleted samples.

**Supplementary Figure 4:** A comparison of the dynamic range for the 4 different technologies using 4 different genes that represent highly expressed (B2M), medium high expression (ACTB), medium low (PGK1) and low (CTCF) expression. These graphs show the expression of the PBMC samples.

**Supplementary Figure 5:** UMAPs cell separations for cell type for each individual technology: 10X 3 prime, 10X FLEX, HIVE and PARSE.

**Supplementary Figure 6:** Bar graphs showing the % neutrophils determined by each technology compared to the flow cytometry result. Each graph displays the data for a single donor.

**Supplementary Figure 7:** Violin plots showing the mitochondrial gene expression levels, n_counts and UMI counts for samples taken 2, 4, 6, 8 and 24 after blood draw. The data was generated using 10X FLEX and each donor is shown individually: A) donor 1, (B) donor 2, (C) donor 3. The % cell type for the time course is shown in (D) donor 1 (E) donor 2 and (F) donor 3.

## Abbreviations

(NETs): Neutrophil Extracellular Traps
(PBMC): Peripheral Blood Mononuclear Cells
(RBC): Red Blood Cell
(single cell RNA-sequencing): scRNA-seq
(UMI): Unique Molecular Identifier

## STAR Methods

Key resources table

Key resources table
| REAGENT or RESOURCE | SOURCE | IDENTIFIER |
| --- | --- | --- |
| **Antibodies** |  |  |
| CD3, BUV805 intracellular | BD Bioscience | AB_2870183 |
| CD4 BUV737 intracellular | BD Bioscience | AB_2870079 |
| CD14 BUV563 surface | BD Bioscience | AB_2870860 |
| CD8 BUV496 surface | BD Bioscience | AB_2870759 |
| CD45 BUV395 surface | BD Bioscience | AB_2869519 |
| HLA-DR BV785 surface | Biolegend | AB_2563461 |
| CD56 BV750 surface | BD Bioscience | AB_2871824 |
| CD11c BV711 surface | BD Bioscience | AB_2738019 |
| CD19 BV650 surface | Biolegend | AB_2564255 |
| CD25 BV605 surface | BD Bioscience | AB_2740127 |
| CD123 BV421 surface | Biolegend | AB_10962571 |
| CD45RA PerCP-Cy5.5 surface | Biolegend | AB_893357 |
| CD15 Pe-Cy7 surface | BD Bioscience | AB_10563901 |
| CD20 Pe-Cy5 surface | Biolegend | AB_314256 |
| CD193 PE-CF594 | BD Bioscience | AB_2737660 |
| FOXP3 PE intracellular | Biolegend | AB_492986 |
| CCR7 APC-Cy7 surface | Biolegend | AB_10916390 |
| CD16 APC surface | Biolegend | AB_314212 |
| human IgG | Jackson ImmunoResearch, West Grove, Pennsylvania | AB_2337043 |
| **Biological samples** |  |  |
| Six healthy volunteer donor blood samples | Roche medical centre |  |
| **Chemicals, peptides, and recombinant proteins** |  |  |
| Phosphate buffered saline (PBS) | ThermoFisher, Massachusetts | Catalogue # 10010023 |
| 2% FBS | VWR International, Pennsylvania | 89510-194 |
| True Stain Monocyte Blocker | Biolegend | 426101 |
| Fixable Green Dead Cell Stain | ThermoFischer Scientific, Waltham, Massachusetts | L23101 |
| Critical commercial assays |  |  |
| Next GEM Single Cell 3<sup>Reagent Kits v3.1</sup> | 10x Genomics, Pleasanton California | PN-1000121 |
| Chromium Next GEM Chip G Single Cell Kit, 16 rxns | 10x Genomics, Pleasanton California | PN-1000127 |
| Single Index Kit T Set A, 96 rxns | 10x Genomics, Pleasanton California | PN-1000213 |
| Chromium Next GEM Single Cell Fixed RNA Sample Preparation Kit | 10x Genomics, Pleasanton California | PN-1000414 |
| Chromium Fixed RNA Kit, Human Transcriptome, | 10x Genomics, Pleasanton California | PN-1000476 |
| Chromium Next GEM Chip Q Single Cell Kit | 10x Genomics, Pleasanton California | PN-1000422 |
| Dual Index Kit TS Set A 96 rxns | 10x Genomics, Pleasanton California | PN-1000251 |
| HIVE collectors | Honeycomb |  |
| HIVE scRNAseq v1 Sample Capture Kit | Honeycomb |  |
| Parse Evercode cell fixation v2 | Parse biosciences |  |
| Evercode™ WT Mini v2 kit | Parse biosciences |  |
| Deposited data |  |  |
| Software and algorithms |  |  |
| Scanpy | https://scanpy.readthedocs.io/en/stable/generated/scanpy.tl.score_genes.html | SCR_018139 |
| Besca | https://github.com/bedapub/besca/blob/master/besca/datasets/genesets/CellNames_scseqCMs6_sigs.gmt |  |
| Uniform Manifold Approximation and Projection (UMAP) algorithm) | https://github.com/lmcinnes/umap | SCR_018217 |
| Leiden algorithm | https://github.com/vtraag/leidenalg |  |
| Msigdb | : https://www.gsea-msigdb.org/gsea/msigdb/human/collections.jsp | SCR_016863 |
| Limma | http://bioinf.wehi.edu.au/limma/ | SCR_010943 |
| Reactome pathways | https://reactome.org | SCR_003485 |
| cellranger 6.0.2 mkfastq | https://www.10xgenomics.com/support/software/cell-ranger/latest/analysis/inputs/cr-mkfastq | SCR_017344 |
| bcl-convert 3.8.4 | https://emea.support.illumina.com/sequencing/sequencing_software/bcl-convert/downloads.html |  |
| split-pipe v1.0.5p: available to PARSE customers on request | https://support.parsebiosciences.com/hc/en-us/articles/20403758539924-How-Do-I-Access-the-Parse-Biosciences-Pipeline |  |
| cellranger | https://www.10xgenomics.com/support/software/cell-ranger/latest/release-notes/cr-release-notes | SCR_017344 |
| Other |  |  |
| 10mL EDTA KCL blood collection tubes |  | 18103 |
| Ficoll | GE healthcare |  |
| SepMate-50 tubes | Stemcell technologies | 85450 |
| 50 ml Falcon tube |  |  |
| EasySep™ RBC Depletion Reagent | Stemcell technologies | 18170 |
| EasySep Easy Eights magnet | Stemcell technologies | 18103 |
| S1 Reagent Kit v1.5 (100 cycles) | Illumina | 20028319 |
| Fixable Green Dead Cell Stain | ThermoFischer Scientific | L23101 |
| 50% Brilliant Stain Buffer | BD Biosciences | 659611 |

## Resource availability

### Lead contact

Further information and requests for resources and reagents should be directed to and will be fulfilled by the lead contact, Emma Bell

### Materials availability

All of the materials used in this study were commercially available

### Data and code availability

• All data reported in this paper will be shared by the lead contact upon request.

• This paper does not report original code

• Any additional information required to reanalyze the data reported in this work paper is available from the lead contact upon request

### Experimental model and study participant details

Whole blood was drawn from 6 healthy human donors. We do not have any information on the sex, gender, ancestry, age, race or ethnicity. These factors are not anticipated to affect the findings of the study which is to compare methods of scRNA-seq as we are not looking at biological variation.

### Method details

#### Samples

Whole blood from healthy donors was drawn into 10mL EDTA KCL or sodium Heparin tubes. For the technology comparison experiments, the blood was divided into 2 ml aliquots, then processed immediately for PBMC isolation or red blood cell removal. In order to determine the time limit for processing and fixing samples after the blood draw we set up a time course. 10 ml of blood was drawn from three donors, a 2 ml aliquot was processed immediately, then subsequent aliquots were processed (RBC depletion) 2, 6, 8 and 24 hours after the original blood draw. In the interim, the sample was stored at 4°C.

#### PBMC Isolation

PBMC were isolated from whole blood using Ficoll (GE healthcare, Illinois) separation according to manufacturer’s instructions. Briefly 15 ml Ficoll was added to SepMate tubes (Stemcell technologies, Vancouver, Canada). Blood was diluted 1:1 with phosphate buffered saline (PBS) (ThermoFischer, Massachusetts) + 2% FBS (VWR International, Pennsylvania).

The diluted blood was then added to the SepMate tube. Tubes were then sealed and centrifuged at 1200xg for 10 mins at room temperature. The top layer of cells, containing enriched mononuclear cells was decanted int,o a new 50 ml Falcon tube. The isolated cells were then washed three times with PBS+2%FBS.

#### RBC Removal

RBCs were removed from blood samples using the EasySep™ RBC Depletion Reagent (Stemcell technologies) and the associated EasySep Easy Eights magnet, according to the manufacturer’s instructions. Briefly, 5mls of whole blood was diluted with PBS-2%FBS solution. The EDTA and RBC depletion solution were added to the samples prior to gently pipette mixing. The samples were transferred to the magnet and left for 5 mins. The supernatant was transferred to a new 14ml tube and the process repeated at least 3 times, or until no RBCs were visibly remaining.

#### Cell counting and viability

After the cell isolations and before fixation or library preparation, the cell viability and number were measured using a Cellaca Cellcounter (Nexcellom Bioscience, Massachusetts). The viability of all samples was high and ranged between 100-96% viability. The number of cells required as input for each different technology is summarized in Table 1. For the library preps for fixed cells, the cell number and viability were counted before fixation and then the fixed cells were counted prior to library prep. For PBMC samples between 250 000-400 000 cells were fixed per sample, and for the RBC depleted samples 500 000-850 000 cells per sample were fixed.

#### 10x 3’transcriptome

An input of 8000 live cells per sample were used for the 10X 3 prime libraries. The libraries were prepared using the Next GEM Single Cell 31 Reagent Kits v3.1. This involves GEM generation on the Chromium instrument, where gel beads were mixed with the live cells and partitioned using oil droplets, resulting in droplets containing a single cell and a gel bead containing barcodes and primers. After barcoding, samples were transferred to a PCR machine for the reverse transcriptase step. Full length, barcoded cDNA from poly adenylated mRNA is then purified using magnetic beads, before PCR amplification. After fragmentation and size selection, P5 and P7 illumina adapters alongside sample barcodes were added to create the final libraries. For sequencing, we targeted 50,000 reads per cell. Therefore, we sequenced the samples on a single NovaSeq S1 flow cell.

#### 10x FLEX

Live cells were fixed using the Chromium Next GEM Single Cell Fixed RNA Sample Preparation Kit (10X Genomics) according to the manufacturer’s instructions. For PBMC samples between 250 000 - 400 000 cells were fixed per sample, and for the RBC depleted samples 500 000-850 000 cells per sample were fixed. Briefly, cells were centrifuged and supernatant removed. Cells were resuspended in the kit supplied fixation buffer, then incubated for 16 hours at 4°C. The cells were centrifuged and supernatant removed before resuspension in the Quenching buffer. The cells were washed, counted and the number of cells adjusted to that of the lowest sample. The Chromium Next GEM Single Cell Fixed RNA Sample Preparation Kit and Chromium Fixed RNA Kit, Human Transcriptome, Chromium Next GEM Chip Q Single Cell Kit were used according to manufacturer’s instructions.

In summary, samples were incubated with the human transcriptome probes to allow hybridization for 16 hours at 42°C. The samples were then washed and transferred to the Chromium X instrument for GEM generation, during this stage the cell is partitioned into an oil droplet containing a 10x barcoded gel bead, so that cell specific barcodes and UMIs are added to the hybridized probes. The probes are then extended and then PCR amplified prior to the addition of P5 and P7 illumina adapters and sample index barcodes. The samples were sequenced on a Novaseq 6000 to a depth of at least 15 000 reads per cell using an SP - 100cycles v1.5 reagent kit.

#### HIVE

Eight HIVE collectors were loaded with freshly isolated cells. HIVE collectors contain more than 65,000 60µm-wide picowells that are pre-loaded with barcoded 3’ transcript capture beads. Each collector was loaded by centrifugation with a total of 15,000 cells according to HIVE scRNAseq v1 Sample Capture Kit User Protocol (Revision A). Once loaded, 3 HIVE devices were incubated for 30 minutes at room temperature before direct processing, and the remaining HIVE devices were frozen at -20°C for later processing after 1 week or 3 weeks of storage. Upon thawing the HIVE devices were equilibrated for 60 minutes at room temperature before processing. All Hive devices, whether processed directly or after storage at -20°C were processed the same way and according to the manufacturer’s instructions.

Briefly, the cells were lysed and hybridized to beads in the HIVE collectors. Beads were recovered by centrifugation into a bead collector and transferred to a filter plate set on a vacuum manifold allowing beads washing and buffer exchanges by aspiration. After the first strand synthesis (45 min at 37°C), 1X NaOH was added and beads washed 3 times before the 2nd strand synthesis (37°C for 30 min). Samples were washed and transferred to a deep well plate for whole transcriptome PCR amplification. After a double-sided SPRI clean-up, samples were indexed by PCR and a final SPRI was performed. Libraries size was checked the bioanalyzer on high sensitivity DNA chips (Agilent), concentration determined on Qubit 3.0 using the dsDNA High sensitivity kit and sequenced on the Novaseq 6000 with a S1 - 100cycles v1.5 reagent kit, using HIVE custom primers and 25-8-8-50 sequencing cycles.

#### PARSE Evercode

Live cells were fixed using the Evercode Cell Fixation v2 kit (Parse Biosciences, Washington). For PBMC samples between 250 000-400 000 cells were fixed per sample, and for the RBC depleted samples 500 000-850 000 cells per sample were fixed according to the manufacturer’s instructions. Cells were spun down at 200 x g at 4°C for 10 mins before resuspension in a prefixation buffer. The cells suspension was then passed through a 40 μm strainer to remove cell clumps. A fixation additive was then added to the suspension and the sample was placed on ice for 10 mins, before the addition of a permeabilizing solution and a further 3 minute incubation on ice. A neutralization buffer was added before a further centrifugation step for 10 minutes, 200 x g at 4°C. The supernatant was removed and cells were resuspended in Cell Buffer and DMSO added to the samples. The fixed cells were then processed using the Evercode™ WT Mini v2 kit according to the manufacturer’s instructions. This protocol uses successive pooling and barcode steps. In the first step, well barcodes are added and reverse transcription of mRNA takes place within the cell. After this step the samples are then pooled and redistributed and further barcodes ligated a further two times. In the third step UMI’s are added to the cDNA. In the last step, cells are lysed, the cDNA isolated and sequencing adapters and sub-library barcodes added by PCR. The resulting libraries were sequenced on the Nova-seq 6000 using an S1 flow cell and S1 Reagent Kit v1.5 (100 cycles) with PE 74bp_6 index_86bp reads

#### Flow Cytometry

250 000 live cells were stained for flow cytometry, cells were centrifuged at 320 x g for 5 mins at 4°C. The supernatant was removed and cells resuspended in 50μl of blocking solution containing 1:100 dilution of human IgG (Jackson ImmunoResearch,West Grove, Pennsylvania), 1:50 dilution True Stain Monocyte Blocker (Biolegend, California) and 1:800 dilution of Fixable Green Dead Cell Stain (ThermoFischer Scientific, Waltham, Massachusetts) in PBS. The samples were incubated at 4°C for 20 minutes. The samples were centrifuged at 320 x g and washed with FACS buffer (PBS supplemented with 2% FCS and 2mM EDTA). The cells were resuspended in 50uL of a solution containing a panel of antibodies for the following surface markers: CD14, CD8, CD45, HLA-DR, CD56, CD11c, CD19, CD25, CD123, CD45RA, LD, CD15, CD20, CD193, CCR7 and CD16. The mix was prepared in a buffer containing 50% FACS buffer and 50% Brilliant Stain Buffer (BD Biosciences, Franklin lakes, New Jersey). The antibodies (previously titrated) are summarized in Table 3. The cells were then incubated for 20 mins at 4°C. Then centrifuged and washed with FACS buffer before incubating for 30 mins with Foxp3 Fixation/Permeabilization working solution (Thermofischer Scientific) at room temperature in the dark. The cells were washed in permeabilization buffer, before the addition of the intracellular antibodies: CD3, CD4 and FOXP3 prepared in permeabilization buffer / Brilliant Stain Buffer (1:1 vol/vol) (details in Supplementary Table 3) and incubated for 30 mins at room temperature. The samples were further centrifuged at 620 x g and resuspended in FACS buffer before running on a FACSymphony™ A5 Cell Analyzer (BD Bioscience).

### Quantification and statistical analysis

#### Bioinformatic Analysis

In total, we sequence 29 samples for the technology comparison, six using Parse, six using 10X Flex, six using 10X 3’, eight using HIVE, and three using 10X 3’ from the HIVE donors. The reads from all technologies were mapped to the human genome (hg38) and we created one gene-by-cell count matrix per sample. FASTQ files from Parse were generated using *bcl-convert 3.8.4* and the *split-pipe v1.0.5p* was utilized to generate the count matrices. FASTQ files from 10X Flex were generated using 10X Genomics *cellranger 6.0.2 mkfastq* and *cellranger 7.1.0 multi* was utilized to generate the count matrices. FASTQ files from 10X 3’ were generated using 10X Genomics *cellranger 6.0.2 mkfastq* and *cellranger 6.0.2 count* was utilized to generate the count matrices. FASTQ files from HIVE were generated using Bcl2fastq for demultiplexing and BeeNet (Honeycomb) for data pre-processing to generate the count matrices.

The count matrices were further processed using BESCA (Madler et al., 2021) and Scanpy (Wolf et al., 2018). Low cutoffs for number of genes expressed and number of UMIs per cell were used in order to capture the neutrophils. This method has been used by previous studies (Wauters et al., 2021), as neutrophils typically have a lower levels of gene expression, and generally have low levels of RNA meaning that they would be filtered out using strict thresholds that have been developed for PBMCs. We applied the same cut-offs for number of genes and UMIs detected for all samples in order to achieve high comparability. Cells that expressed at least 200 and not more than 10,000 genes and included at least 200 and not more than 50,000 UMIs were kept for downstream analysis. In addition, we removed cells with high mitochondrial gene expression, for Parse PBMC samples above 1%, Parse RBC depleted samples above 2%, and all other samples above 10% of UMIs mapping to mitochondrial genes. This resulted in 332,783 total cells.

Normalization was performed for the entire dataset using count depth scaling to 10,000 total counts per cell, resulting in the cp10k (counts per 10,000) unit. Count values were log-transformed using natural logarithm: ln(cp10k1+11). To reduce dataset dimensionality before clustering, the highly variable genes within the dataset were selected. Genes were defined as being highly variable when they have a minimum mean expression of 0.0125, a maximum mean expression of 3 and a minimum dispersion of 0.5. Technical variance was removed by regressing out the effects of count depth and mitochondrial gene content and the gene expression values were scaled to a mean of 0 and variance of 1 with a maximum value of 10. The first 50 principal components were calculated and used as input for calculation of the 10 nearest neighbours. The neighbourhood graph was then embedded into two-dimensional space using the Uniform Manifold Approximation and Projection (UMAP) algorithm) [25] Cell communities are detected using the Leiden algorithm (Traag et al., 2019) at a resolution of 1.

We performed cell type annotation for the clustering of cells from each technology individually and mapped it to the entire dataset. We assessed the cell types by calculating signature scores for all signatures provided by Besca (https://github.com/bedapub/besca/blob/master/besca/datasets/genesets/CellNames_scse <u>qCMs6_sigs.gmt</u>). The score is the average expression of a set of genes subtracted with the average expression of a reference set of genes, calculated by Scanpy’s *score_genes* function (https://scanpy.readthedocs.io/en/stable/generated/scanpy.tl.score_genes.html). For the cell type composition comparison we removed clusters of doublets and other artifacts before calculating cell type fractions per sample.

For each single cell sequencing technology, we calculated the mean absolute error (MAE) and root mean squared error (RMSE) of the neutrophil abundance compared to the abundance obtained by FACS. For each donor, we averaged the neutrophil abundance from three FACS experiments (technical replicates). Then, for each donor and technology, we determined the absolute difference between the FACS-derived abundance and the sequencing technology-derived abundance. The average of these differences across donors is the mean absolute error (MAE) for each technology. Similarly, we calculated the root mean squared error (RMSE) by first determining the squared differences between the FACS-derived and sequencing technology-derived abundances, then averaging these squared differences, and finally taking the square-root of this average for each technology.

For the time course experiment, we sequence 18 samples, all on the 10X Flex technology. The dataset was processed as described above, but different filtering criteria were applied to the cells: number of genes: 200 - 4,000; number of UMI counts 200 - 15,000; maximal mitochondrial fraction 1% (see Suppl. Fig. 7a-c). In order to identify differentially expressed genes in neutrophils over time, we generated a pseudobulk gene-by-sample matrix for the neutrophils identified. We tested each time point (2h, 4h, 6h, 8h, 24h) versus immediate processing (0h) by fitting the model: ∼ 0 + Donor + Time point. We filtered genes at a cut-off of average transcripts per million larger than 0.25. 12,726 genes were assessed for differential expression using limma+voom (source: http://bioinf.wehi.edu.au/limma/). We chose relaxed cut-offs to allow for high sensitivity to detect changes, fold-change larger than 1.5 or less than 0.666 and false discovery rate smaller than 10%. Afterwards we counted the number of up- or down-regulated genes according to these cut-offs. We applied gene set enrichment analysis using Camera (Wu and Smyth, 2012). We used multiple geneset collections, including MSigDB hallmark (source: https://www.gsea-msigdb.org/gsea/msigdb/human/collections.jsp), Reactome pathways (source: https://reactome.org), and an internally curated set of cancer immuno-therapy (CIT) signatures. The latter includes a Stress MP5 (meta-program 5) signature discovered by (Gavish et al., 2023) which showed highest enrichment.

