## Supplemental Tables for "Comparison of Fixed Single Cell RNA-seq Methods to Enable Transcriptome Profiling of Neutrophils in Clinical Samples"

**Table 1**

| **Method** | **Input** | **Barcoding strategy** | **Throughput** | **Description** | **Storage and shipment** |
| --- | --- | --- | --- | --- | --- |
| 10X Genomics: 10x 3 prime transcriptome | Live cells | Cells are captured in GEMs for barcoding and UMI on transcript | Maximum of 10,000 cells (increasing cell number increases the doublet rate) | Single cells are partitioned into oil droplets containing a gel bead with necessary barcodes | Well established protocols for PBMCs (freeze cells down and gently defrost before processing)  Not suitable for granulocytes |
| 10X Genomics: 10X FLEX RNA | Fixed cells | Probes are hybridised to mRNA in fixed cells prior to partitioning in oil droplets with a gel bead containing barcodes | Recommended collection of ≥300,000 cells or ≥500,000 nuclei per sample | Cells are fixed at the point of collection with paraformaldehyde. Flex works by using hybridisation to mRNA transcripts, barcoded probes are sequenced | Fixed cells are suitable for freezing and shipping. Stability confirmed up to 6 months |
| Parse biotechnologies Evercode | Fixed cells | A combinatorial barcoding strategy is used on fixed cells. The cell acts as a partition where reverse transcription and barcode ligation steps take place | Flexible cell number input 1000-1 million cells | No specialist instruments required  Better at detecting low expressed mRNAs | Fixed cells are suitable for freezing and shipping. |
| Honeycomb, HIVE | Live cells (frozen) | Barcoded mRNA-capture beads: 1 bead one cell | Recommended loading 15000 cells (v1) | Cells settle into picowells containing barcoded mRNA-capture beads. Lyse cells in HIVE device and perform library prep. No specialist instruments required. | HIVE device can be stored in the freezer or shipped to analytical labs |

**Table 2**

| **Technology** | **PBMC** | | | **RBC depleted** | | |
| --- | --- | --- | --- | --- | --- | --- |
|  | **Mean (%)** | **MAE** | **RMSE** | **Mean**  **(%)** | **MAE** | **RMSE** |
| 10x 3 prime transcriptome | 0.00 | 0.20 | 0.25 | 0.00 | 38.13 | 38.33 |
| 10X FLEX RNA | 0.54 | 0.34 | 0.39 | 44.31 | 6.18 | 6.41 |
| Parse biotechnologies Evercode | 0.91 | 0.71 | 0.85 | 22.41 | 15.73 | 20.93 |
| Honeycomb, HIVE | - | - | - | 40.92 | 17.38 | 17.95 |

**Supplementary Table 1**

| **Cell type** | **% cell population** | | | | | | | |
| --- | --- | --- | --- | --- | --- | --- | --- | --- |
|  | **Flow cytometry**  **RBC depleted** | **Flow cytometry PBMC** | **10x 3 prime**  **RBC depleted** | **10x 3 prime**  **PBMC** | **Flex**  **RBC depleted** | **Flex**  **PBMC** | **PARSE**  **RBC depleted** | **PARSE**  **PBMC** |
| B cell | 3.43 | 5.69 | 4.49 | 4.76 | 3.86 | 5.78 | 5.96 | 6.28 |
| T cell | 38.7 | 69.1 | - | - | 37.38 | 66.14 | 36.9 | 67.74 |
| Monocyte | 4.05 | 2.92 | 7.57 | 3.51 | 5.07 | 2.9 | 7.97 | 2.52 |
| Natural Killer cells | 10.8 | 17.3 | - | - | 13.77 | 24.89 | 15.33 | 18.79 |
| Neutrophils | 34.5 | 0.044 | 0.0 | 0.0 | 39.23 | 0.18 | 29.11 | 0.26 |

**Supplementary Table 2 (Donor 2)**

| **Cell type** | **Flow cytometry**  **RBC depleted** | **Flow cytometry PBMC** | **10x 3 prime**  **RBC depleted** | **10x 3 prime**  **PBMC** | **Flex**  **RBC depleted** | **Flex**  **PBMC** | **PARSE**  **RBC depleted** | **PARSE**  **PBMC** |
| --- | --- | --- | --- | --- | --- | --- | --- | --- |
| B cell | 6.91 | 12.0 | 13.17 | 13.58 | 7.5 | 14.6 | 14.19 | 14.37 |
| T cell | 37.0 | 67.1 | - | - | 31.68 | 67.01 | 28.07 | 61.98 |
| Monocyte | 6.2 | 7.42 | 9.75 | 5.77 | 7.32 | 6.57 | 16.72 | 5.7 |
| Natural Killer cells | 5.0 | 8.41 | - | - | 5.23 | 10.58 | 6.42 | 8.08 |
| Neutrophils | 36.4 | 0.39 | 0.0 | 0.0 | 41.62 | 0.67 | 29.86 | 0.95 |

**Supplementary Table 3 (Donor 3)**

| **Cell type** | **Flow cytometry**  **RBC depleted** | **Flow cytometry PBMC** | **10x 3 prime**  **RBC depleted** | **10x 3 prime**  **PBMC** | **Flex**  **RBC depleted** | **Flex**  **PBMC** | **PARSE**  **RBC depleted** | **PARSE**  **PBMC** |
| --- | --- | --- | --- | --- | --- | --- | --- | --- |
| B cell | 3.28 | 7.13 | 7.0 | 9.58 | 3.73 | 8.41 | 13.16 | 8.99 |
| T cell | 27.1 | 60.8 | - | - | 24.62 | 66.14 | 24.06 | 55.27 |
| Monocyte | 9.91 | 15.8 | 27.45 | 14.19 | 11.02 | 11.06 | 44.07 | 21.1 |
| Natural Killer cells | 4.23 | 8.16 | - | - | 4.58 | 10.9 | 7.18 | 7.41 |
| Neutrophils | 43.5 | 0.17 | 0.0 | 0.0 | 52.09 | 0.77 | 8.25 | 1.51 |

**Supplementary Table 4 (Donor 4)**

| **Cell type** | **Flow cytometry**  **RBC depleted** | **HIVE**  **RBC depleted** |
| --- | --- | --- |
| B cell | 2.64 | 1.71 |
| T cell | 16.4 | 19.42 |
| Monocyte | 9.16 | 23.51 |
| Natural Killer cells |  |  |
| Neutrophils | 63.3 | 51.79 |

**Supplementary table 5: Donor 5**

| **Cell type** | **Flow cytometry**  **RBC depleted** | **HIVE**  **RBC depleted** |
| --- | --- | --- |
| B cell | 1.78 | 2.82 |
| T cell | 21.1 | 31.51 |
| Monocyte | 6.26 | 21.39 |
| Natural Killer cells |  |  |
| Neutrophils | 63.6 | 41.23 |

**Supplementary table 6: Donor 6**

| **Cell type** | **Flow cytometry**  **RBC depleted** | **HIVE**  **RBC depleted** |
| --- | --- | --- |
| B cell | 4.57 | 6.33 |
| T cell | 24.2 | 30.11 |
| Monocyte | 11.0 | 29.51 |
| Natural Killer cells |  |  |
| Neutrophils | 48.0 | 29.73 |

**Supplementary Table 7**

| **Target** | **Fluorochrome** | **Intracellular or surface marker** | **Dilution** | **Catalog No** | **Vendor** |
| --- | --- | --- | --- | --- | --- |
| CD3 | BUV805 | intracellular | 400 | 612895 | BD |
| CD4 | BUV737 | intracellular | 50 | 612748 | BD |
| CD14 | BUV563 | surface | 50 | 741360 | BD |
| CD8 | BUV496 | surface | 25 | 741199 | BD |
| CD45 | BUV395 | surface | 100 | 563792 | BD |
| HLA-DR | BV785 | surface | 100 | 307642 | Biolegend |
| CD56 | BV750 | surface | 200 | 747068 | BD |
| CD11c | BV711 | surface | 50 | 563130 | BD |
| CD19 | BV650 | surface | 100 | 363026 | Biolegend |
| CD25 | BV605 | surface | 50 | 740397 | BD |
| CD123 | BV421 | surface | 50 | 306018 | Biolegend |
| CD45RA | PerCP-Cy5.5 | surface | 400 | 304122 | Biolegend |
| Live/Dead | GREEN | surface | 800 | L34970 | Thermofisher |
| CD15 | Pe-Cy7 | surface | 100 | 560827 | BD |
| CD20 | Pe-Cy5 | surface | 100 | 302308 | Biolegend |
| CD193 | PE-CF594 | surface | 100 | 562571 | BD |
| FOXP3 | PE | intracellular | 50 | 320108 | Biolegend |
| CCR7 | APC-Cy7 | surface | 200 | 353212 | Biolegend |
| CD16 | APC | surface | 200 | 302012 | Biolegend |
