## Supplemental Figures for "Comparison of Fixed Single Cell RNA-seq Methods to Enable Transcriptome Profiling of Neutrophils in Clinical Samples"

### Slide 1
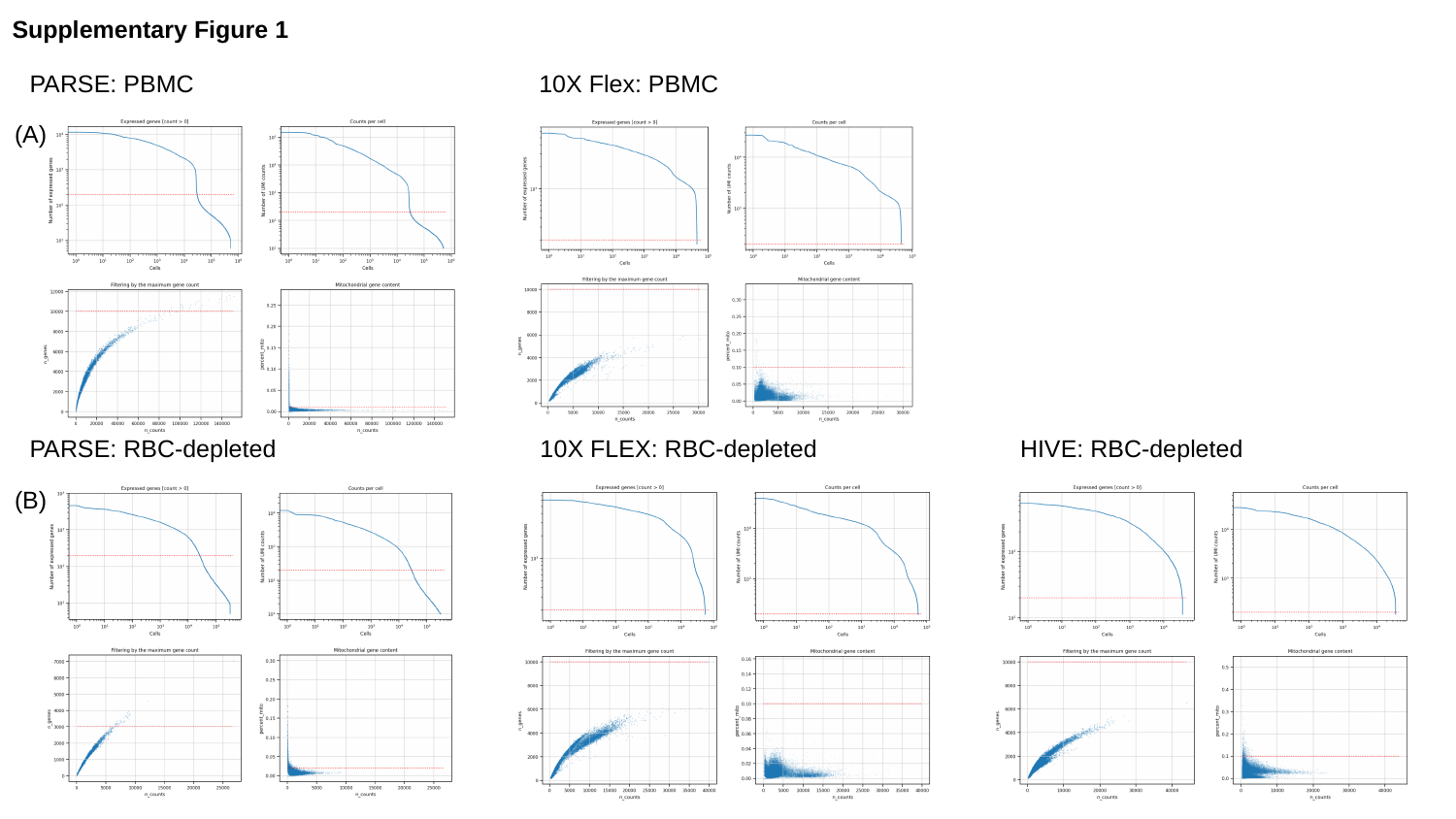

Supplementary Figure 1
PARSE: PBMC 10X Flex: PBMC
(A)
PARSE: RBC-depleted 10X FLEX: RBC-depleted HIVE: RBC-depleted
(B)

### Slide 2
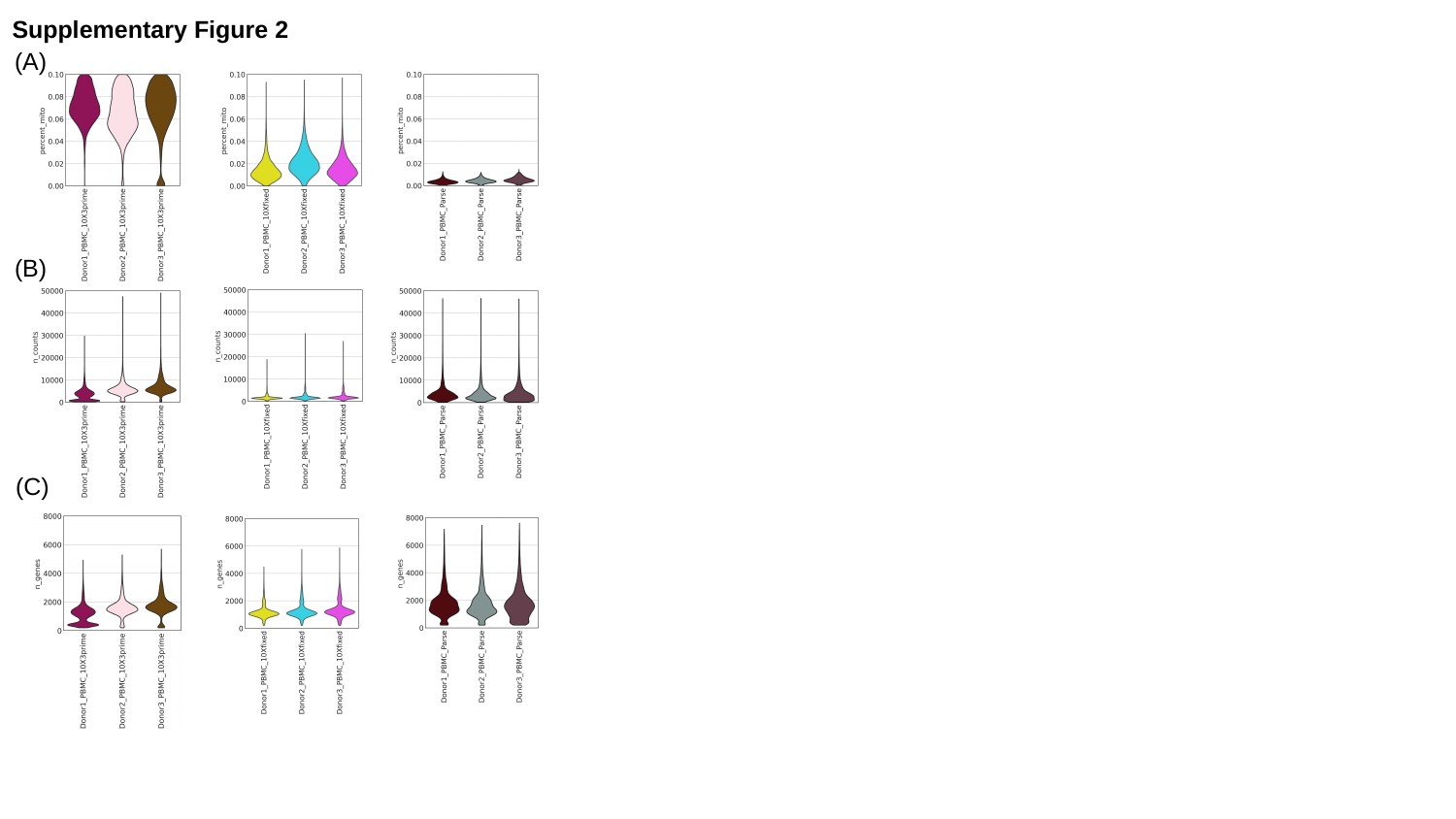

Supplementary Figure 2
(A)
(B)
(C)

### Slide 3
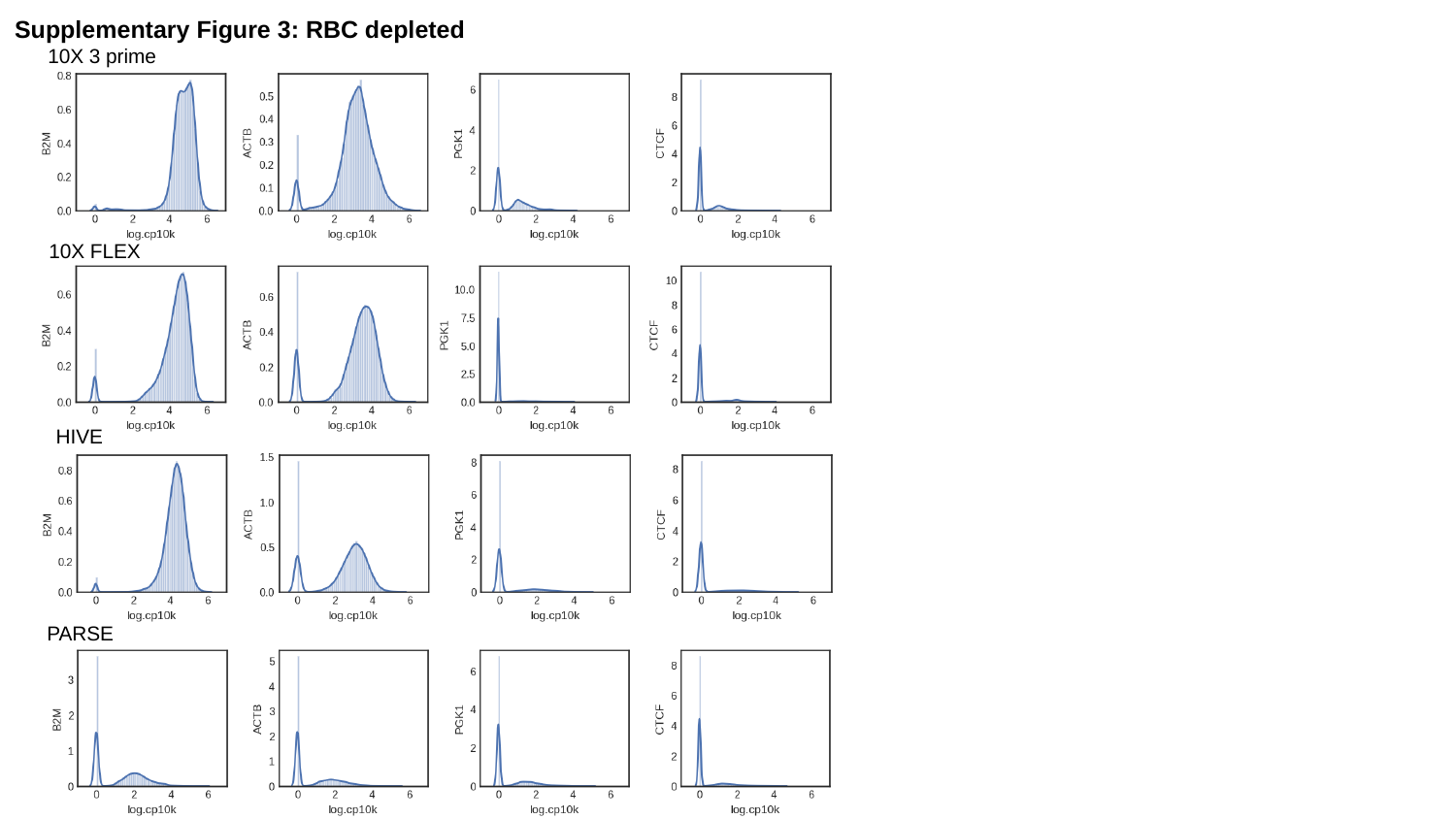

Supplementary Figure 3: RBC depleted
10X 3 prime
10X FLEX
HIVE
PARSE

### Slide 4
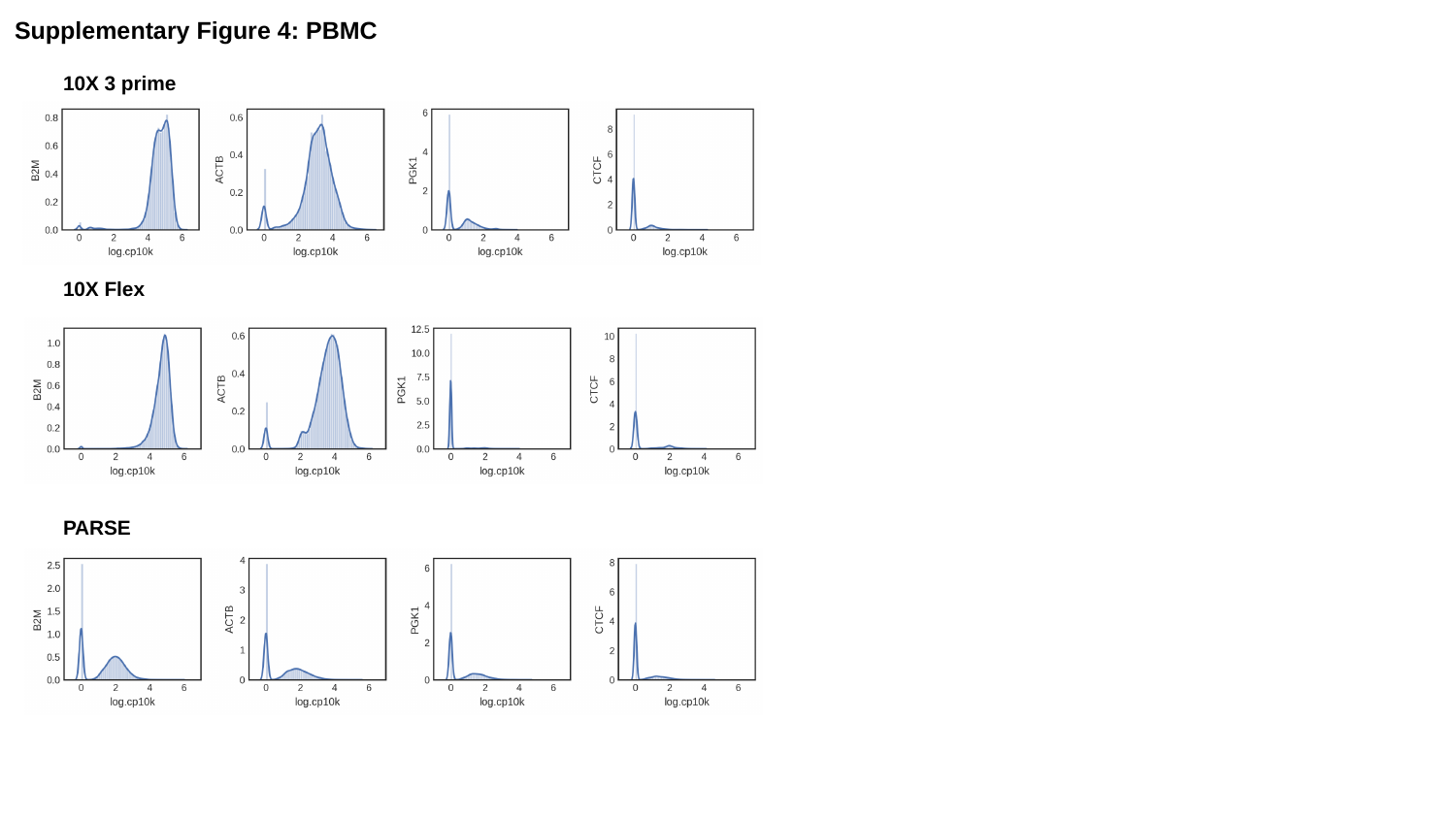

Supplementary Figure 4: PBMC
10X 3 prime
10X Flex
PARSE

### Slide 5
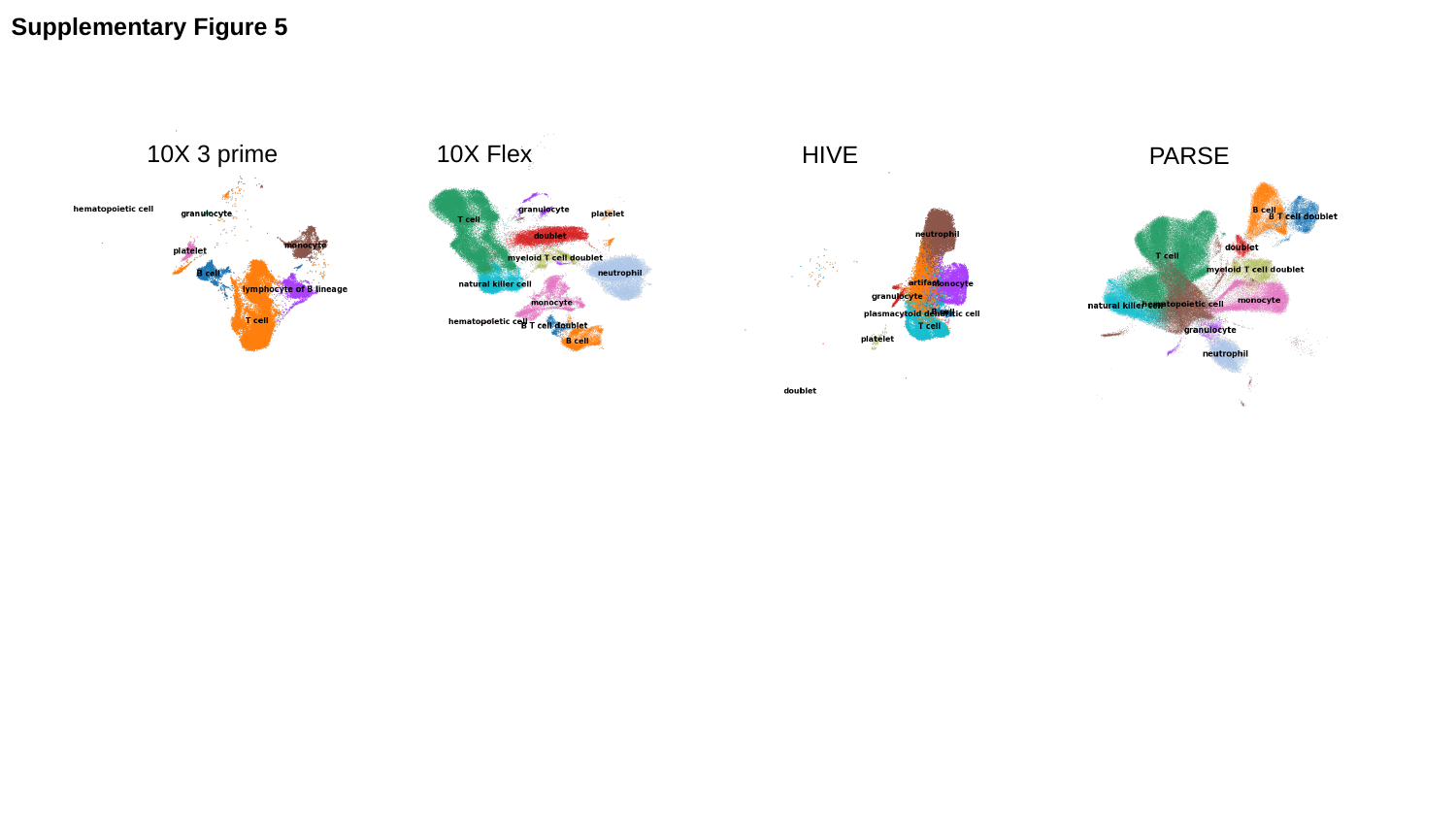

Supplementary Figure 5
10X 3 prime
10X Flex
HIVE
PARSE

### Slide 6
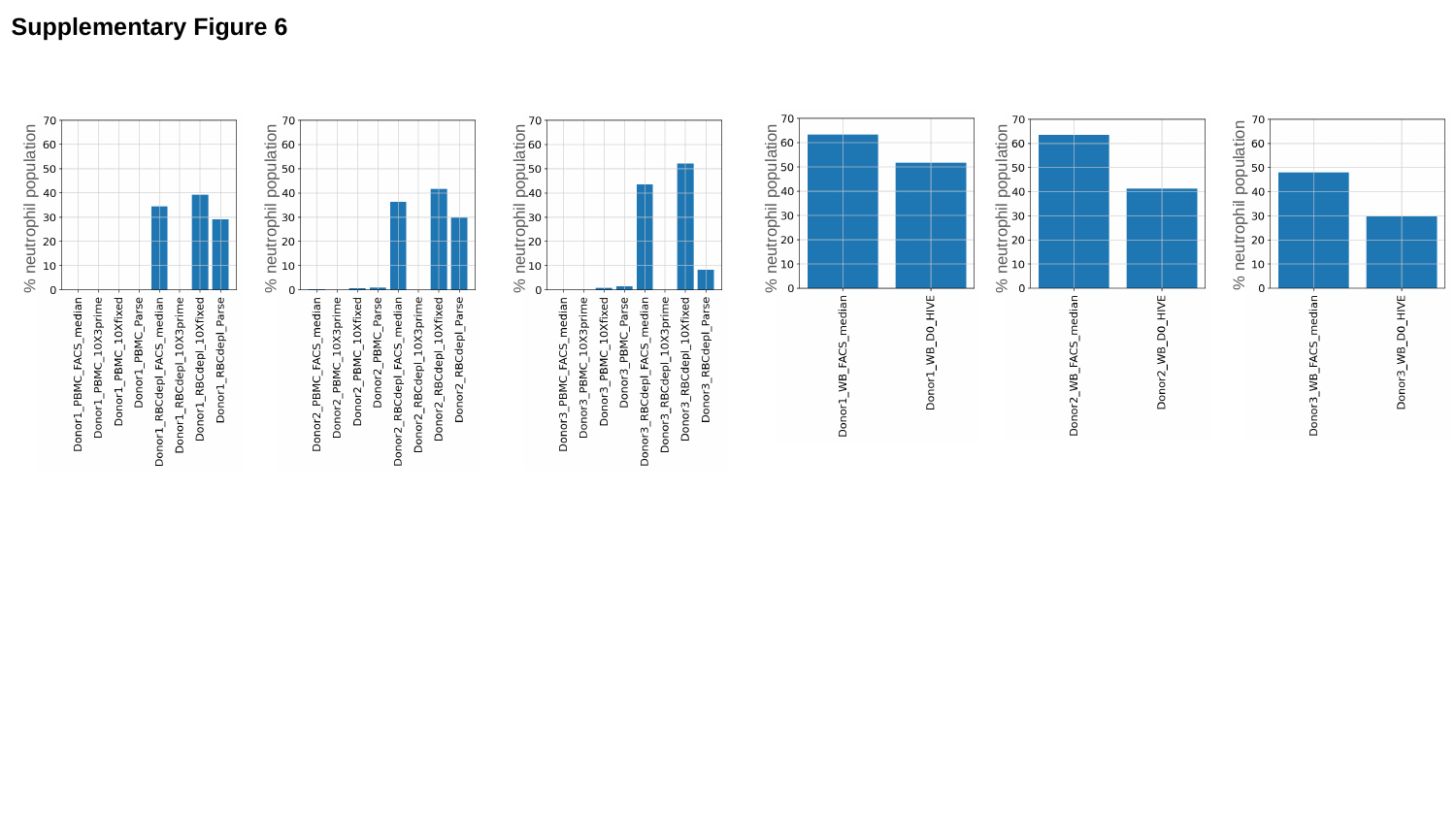

Supplementary Figure 6
% neutrophil population
% neutrophil population
% neutrophil population
% neutrophil population
% neutrophil population
% neutrophil population

### Slide 7
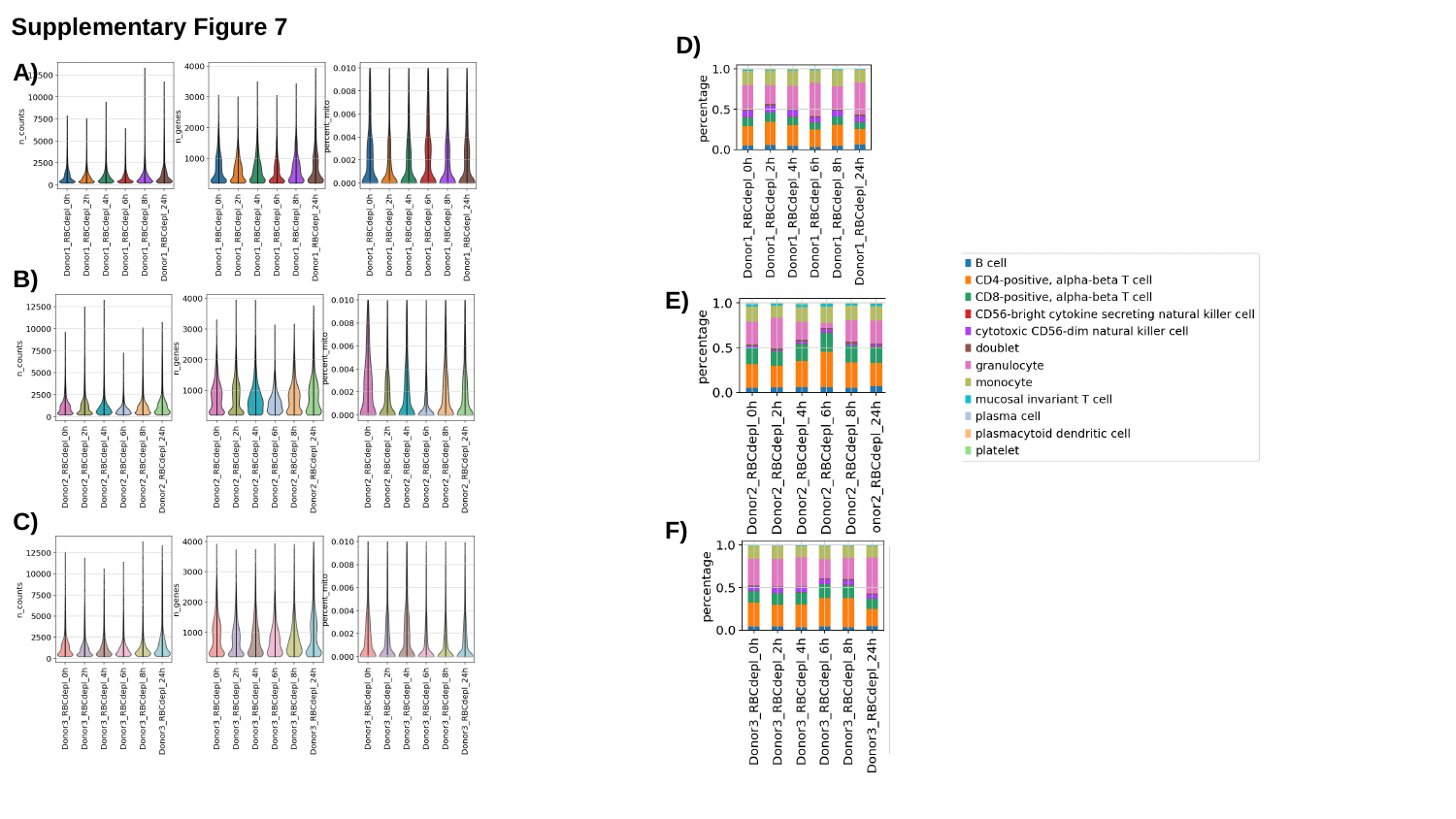

Supplementary Figure 7
D)
A)
B)
E)
C)
F)
